# Investigation of the futalosine pathway for menaquinone biosynthesis as a novel target in the inhibition of *Chlamydia trachomatis* infection

**DOI:** 10.1101/2021.10.26.465979

**Authors:** Brianne M. Dudiak, Tri M. Nguyen, David Needham, Taylor C. Outlaw, Dewey G. McCafferty

## Abstract

*Chlamydia trachomatis,* an obligate intracellular bacterium with limited metabolic capabilities, possesses the futalosine pathway for menaquinone biosynthesis. Futalosine pathway enzymes have promise as narrow spectrum targets, but the activity and essentiality of chlamydial menaquinone biosynthesis have yet to be established. In this work, menaquinone-7 (MK-7) was identified as a *C. trachomatis*-produced quinone through LC-MS/MS. An immunofluorescence-based assay revealed that treatment of *C. trachomatis*-infected HeLa cells with futalosine pathway inhibitor docosahexaenoic acid (DHA) reduced inclusion number, inclusion size, and infectious progeny. Supplementation with MK-7 nanoparticles rescued the effect of DHA on inclusion number, indicating that the futalosine pathway is a target of DHA in this system. These results open the door for menaquinone biosynthesis inhibitors to be pursued in antichlamydial development.

## INTRODUCTION

*Chlamydia trachomatis* is the pathogen responsible for the most predominant bacterial sexually transmitted infection worldwide, with over 130 million new infections reported each year.^1,2^ Because over 70 percent of chlamydial infections are initially asymptomatic, infections are often undiagnosed.^3^ Left untreated, chlamydial infections can lead to the development of severe sequelae such as pelvic inflammatory disease, infertility, and ectopic pregnancy,^4^ as well as increased susceptibility to HIV infection and cervical cancer.^5,6^

The current therapeutics of choice for chlamydial infections are the broad-spectrum antibiotics azithromycin and doxycycline.^7^ Despite the effectiveness of these front-line antibiotics to date, chlamydial infections still place an extraordinary burden on public health. Complex^8^ and partial^9^ protective immunity leads to frequent re-infections. If *C. trachomatis* is exposed to stressors such as β-lactam antibiotics, nutrient deprivation, or human cytokines,^10^ the bacterium enters a temporary, aberrant phase called persistence, which is characterized by metabolic quiescence and an inability to replicate until the stressor has been removed.^11^ Significantly, persistent *C. trachomatis* has been shown to be less susceptible to azithromycin in a mouse model of chlamydial infection.^12^ Recent reports have also indicated that repeat infections due to azithromycin treatment failure are an emerging concern.^13,14^ Collectively, the overwhelming number of cases per year in combination with the increase in infection reoccurrences underscores the need to investigate the complex infection biology of *C. trachomatis* and explore new therapeutic strategies to combat this pathogen.

*C. trachomatis* is an obligate intracellular bacterium with a unique biphasic life cycle comprised of the infectious elementary body phase and the replicative reticulate body phase.^15^ The life cycle begins with internalization of elementary bodies into the outermost layer of epithelial cells of the genital tract or eyes by endocytosis. A parasitophorous vacuole, termed an inclusion, encapsulates the elementary bodies to shield *C. trachomatis* from the host immune response. Within the inclusion, elementary bodies differentiate into reticulate bodies, which replicate exponentially by binary fission. Reticulate bodies then transition back into elementary bodies for release from the host cell by lysis or extrusion and initiation of a new round of infection.

As an obligate intracellular pathogen, *C. trachomatis* relies extensively on host-derived nutrients for survival. *C. trachomatis* has a minimal genome of approximately 900 protein-encoding genes,^16^ leaving the pathogen with limited metabolic capabilities.^17^ As such, *C. trachomatis* was historically believed to be an energy parasite that exclusively imported ATP from the host cell until the recent discovery of a functional sodium-dependent respiratory chain.^18^ For an electron transport chain to produce ATP, a membrane-associated electron carrier is required to shuttle electrons between respiratory complexes.^19^ Interestingly, despite *C. trachomatis* lacking many biosynthetic pathways, the full suite of genes required for production of the electron carrier menaquinone have been identified in the chlamydial genome.^20^ *C. trachomatis* harbors the genes for the futalosine pathway for menaquinone biosynthesis (Figure 1, Figure S1), which is enzymatically distinct from the more prevalent classical pathway.^21^ While menaquinone biosynthesis by *C. trachomatis* has not been confirmed to date, production of menaquinone through the futalosine pathway is experimentally supported by the biophysical and enzymatic characterization of CT263 (MqnB), a 6-amino-6-deoxyfutalosine hydrolase responsible for the third step in the pathway.^22^

**Figure 1.**
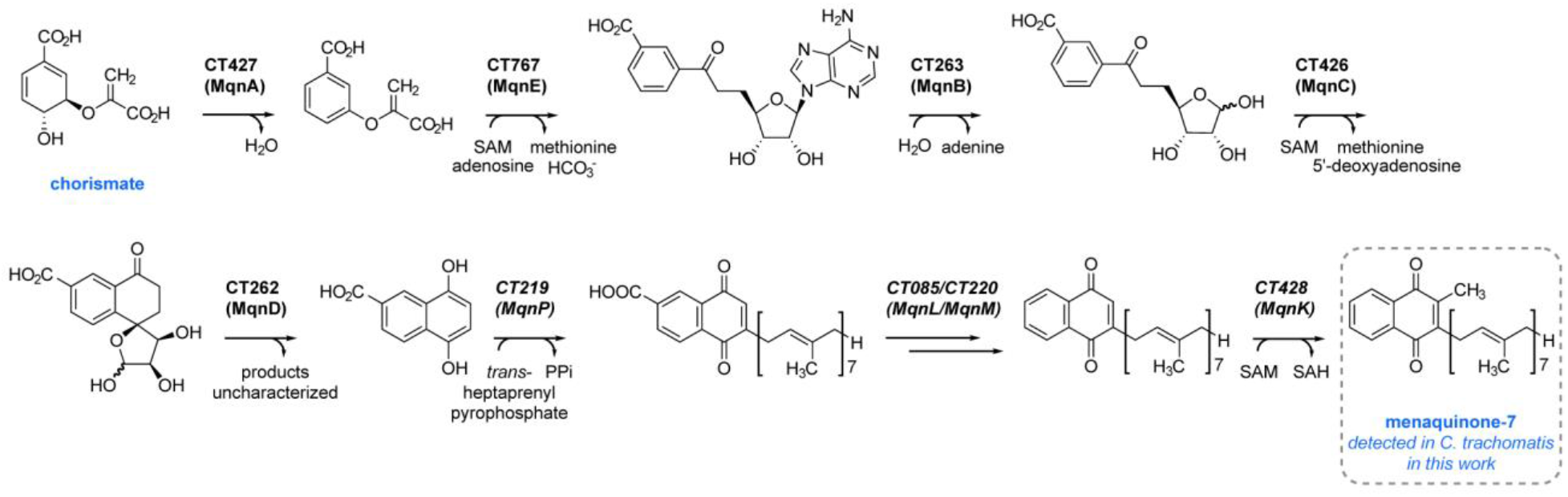
Predicted futalosine pathway in *C. trachomatis.* Menaquinone-7 biosynthesis via the futalosine pathway occurs through the indicated enzymatic steps, with hypothesized chlamydial enzymes annotated. Steps from initial precursor chorismate to 1,4-dihydroxy-6-naphthoic acid have been characterized in *Streptomyces coelicolor.*^25^ The order of the last three steps (italicized) has not been experimentally confirmed (Figure S1).

Targeting the futalosine pathway presents an attractive opportunity for therapeutic intervention. Inhibitors of Na-NQR, the first complex of the chlamydial respiratory chain, have been demonstrated to inhibit chlamydial growth and replication, suggesting that respiratory chain function is important for survival.^18,23^ Further, the futalosine pathway has only been identified in a small group of bacteria and is not present in humans or commensal bacteria.^20^ Humans utilize ubiquinone, a different electron carrier, in the mitochondrial electron transport chain,^24^ while commensal bacteria that produce menaquinone exclusively possess the classical pathway.^25^ As such, the chlamydial futalosine pathway enzymes have promise as potential narrow spectrum targets; however, the effects of disrupting menaquinone biosynthesis in *C. trachomatis* have yet to be elucidated.

We hypothesized that due to the importance of the respiratory chain in *C. trachomatis* and the critical role of an electron carrier in electron transport, menaquinone biosynthesis may be essential for a functional respiratory chain and significant for chlamydial viability. Here we describe mass spectrometry and chemical genetics analyses to characterize the function of the futalosine pathway and investigate the effects of futalosine pathway inhibitors on chlamydial infection. We identified menaquinone-7 (MK-7) as a chlamydial quinone by tandem mass spectrometry, confirming that the futalosine pathway is functional in *C. trachomatis*. We subsequently challenged *C. trachomatis*-infected HeLa cells with inhibitors of the futalosine pathway identified previously in other organisms and found that (4Z,7Z, 10Z,13Z,16Z,19Z)-docosa-4,7,10,13,16,19-hexaenoate (DHA) treatment reduces chlamydial inclusion number, inclusion size, and infectious progeny. The effect of DHA on inclusion number was rescued by concurrent treatment with MK-7 nanoparticles, which is consistent with a mechanism of targeting the futalosine pathway. Together, these results illustrate that menaquinone biosynthesis is a previously unexploited target in *C. trachomatis* that merits further investigation in the development of new antichlamydial agents.

## MATERIALS AND METHODS

### General materials and methods

Chemicals were purchased from Millipore Sigma, Fisher Scientific, VWR, or Cayman Chemicals unless otherwise stated and used without further purification. Mass spectrometry (MS) data were obtained on an Agilent 6460 Triple Quadrupole LC-MS. Absorbance measurements were obtained with a SpectraMax i3X plate reader (Molecular Devices). Representative microscopy images were obtained on a Zeiss Axio Observer widefield fluorescence microscope with a 10X objective and processed with ImageJ.^26^ Data were visualized using GraphPad Prism.

### Chlamydial and mammalian cell culture

*Homo sapiens* HeLa cells (Duke Cell Culture Facility via ATCC, CCL-2) were cultured in Dulbecco’s modified Eagle’s medium (DMEM, ThermoFisher Scientific) supplemented with 10% heat-inactivated fetal bovine serum (FBS, Millipore Sigma) at 37 °C and 5% CO2. *C. trachomatis* L2 (strain 434/Bu) stocks were prepared in sucrose-phosphate-glutamate (SPG) buffer as described previously,^27^ and aliquots were stored at −80 °C until needed for infections.

### Quinone extractions and detection by LC-MS/MS

Isoprenoid quinone-containing samples were prepared based on previously reported methodology for extraction of isoprenoid quinones from plasma.^28^ HeLa cells were trypsinized, centrifuged at 180 x *g* for 4 min, washed with PBS, centrifuged again, and re-suspended in ice-cold ethanol. *C. trachomatis* stocks were thawed, centrifuged at 18,000 x *g* for 30 minutes, and resuspended in ice-cold ethanol. Both samples were sonicated (four 15 s pulses at 30% amplitude) and subjected to three rounds of extractions. In each extraction, hexanes was added, the solution was vortexed and centrifuged at 1,400 x *g* for 5 min, and the hexanes layer was reserved.^28^ Hexanes layers for each sample were combined and concentrated to 400 μL under a stream of N2 (g). An equal volume of isopropanol was added to each sample, and the mixtures were incubated at −20 °C for 40 min. Samples were centrifuged at 3,000 x *g* for 10 min to pellet precipitate, and the supernatants were removed and concentrated to 250 μL. The incubation and centrifugation steps were repeated, and samples were concentrated to dryness. Residues were dissolved in methanol for LC-MS/MS analysis. A 100 ng/mL solution of menaquinone-7 in methanol was prepared as a standard.

Samples were analyzed by LC-MS/MS with an XDB-C8, 2.1 x 50 mm, 3.5 μm column (Agilent Zorbax Eclipse). A 2 min isocratic method with a mobile phase of 5 mM ammonium formate in methanol was employed. Precursor ion scans for the daughter ion m/z 187.0 were conducted in positive ion mode with a fragmentor voltage of 185 V and collision energy of 30. Extracted ion chromatograms (EICs) were collected for the [M+H]^+^ MK-7 species.

### Chlamydial inclusion experiments

Immunofluorescence microscopy experiments for evaluating the effects of antichlamydial agents were conducted based on previous reports.^29,30^ HeLa cells seeded in 96 well plates were grown to 90-100% confluency, then infected with *C. trachomatis* at an MOI of 0.5. Infections were facilitated by centrifugation at 900 x *g* for 1 hr. Following centrifugation, the culture media was replaced with fresh media containing an inhibitor of interest or a vehicle control. After 40 hours of incubation at 37 °C and 5% CO2, cells were fixed with ice-cold methanol for visualizing inclusions. Methanol-fixed cells were stained for immunofluorescence microscopy, with either a fluorescein-conjugated chlamydial lipopolysaccharide antibody (Pathfinder Chlamydia Confirmation System, BioRad) or a major outer membrane protein (MOMP) antibody (1:1000, Novus Bio) and an appropriate AlexaFluor secondary antibody (1:200, ThermoFisher Scientific). Counterstaining was performed with Hoechst stain. Images of 9 fields per well were obtained using the CellInsight CX5 High-Content Screening Platform (ThermoFisher Scientific) with a 10X objective. Inclusion number and size were measured using HCS Studio Cell Analysis Software. Data are shown as the average inclusion number or size for a treatment condition relative to that of the respective ethanol (12-methyltetradecanoic acid and DHA) or DMSO (BuT-DADMe-ImmA and siamycin I) vehicle control ± s.e.m.

### Infectious progeny experiments

*C. trachomatis* infections with inhibitor treatment were performed as described above, but in 24 well plates. At 40 hours postinfection (hpi), infected cells were isolated by scraping and collecting in the original culture media and were stored at −80 °C until analysis by titer assay. Samples were thawed, vortexed, and serially diluted (1:10). Diluted samples were used to infect new HeLa cells seeded in 96 well plates. At 40 hpi, the previously described immunofluorescence staining and imaging protocol was employed to count recoverable IFUs. Data are shown as the average infectious progeny for a treatment condition relative to that of the ethanol vehicle control ± s.e.m.

### Menaquinone-7 nanoparticle preparation

Menaquinone-7 nanoparticles were prepared using a rapid solvent exchange method.^31^ Briefly, a solution of menaquinone-7 in ethanol (500 μL) was rapidly injected into 4.5 mL of ultrapure water stirring at 500 rpm, resulting in the spontaneous generation of nanoparticles. Nanoparticles were sterile-filtered and measured using a dynamic light scattering instrument (Zetasizer, Malvern).

### MTT viability assay

HeLa cells seeded in 96 well plates were treated with various concentrations of DHA or the corresponding percentage of ethanol in DMEM. At 40 hours post-treatment, media was removed from each well and replaced with equal volumes of DMEM and a 5 mg/mL solution of MTT in PBS.^32^ The plates were incubated at 37 °C for 3 h. Formazan crystals were solubilized by addition of 4 mM HCl, 0.1% NP-40 in isopropanol, and the plate was placed on an orbital shaker for 15 min. Absorbances were measured at 570 nm using a plate reader and are represented as the normalized absorbance relative to that of untreated HeLa cells ± s.e.m.

### Statistical analysis

Statistical analyses were performed using JMP.^33^ For experiments determining the effects of DHA on inclusion number, inclusion size, and infectious progeny, the raw data from three independent experiments were evaluated by RM-ANOVA with Tukey’s post hoc test for pairwise comparisons, where p < 0.05 was considered statistically significant. For ease of visualization, data are shown as percentages of the respective vehicle control values.

## RESULTS AND DISCUSSION

### Menaquinone-7 is a chlamydial isoprenoid quinone

To validate the activity of the futalosine pathway in *C. trachomatis*, we performed a liquid chromatography-coupled tandem mass spectrometry (LC-MS/MS) analysis of chlamydial and uninfected HeLa cell extracts to identify a chlamydia-produced menaquinone. The uninfected HeLa cell sample was included as a negative control, as the *H. sapiens* cervical cell line does not possess the futalosine pathway. Chlamydial and uninfected HeLa cell samples were first subjected to hexanes extractions to isolate isoprenoid quinones.^28^ Because bacterial menaquinones have polyisoprenyl side chains of varying lengths,^34^ we employed an

LC-MS/MS precursor ion scan for the 2-methyl-1,4-naphthoquinone product ion (m/z 187.0).^35^ Analysis of both extraction samples identified MK-7 exclusively in the chlamydial extract as the [M+H]^+^ and [M+NH_4_]^+^ ions (observed [M+H]^+^=649.4, calculated [M+H]^+^=649.5; observed [M+NH_4_]^+^=666.3, calculated [M+NH_4_]^+^=666.5) (Figure 2). MK-7 extracted from *C. trachomatis* had the same retention time as a commercially obtained MK-7 standard, further verifying the identity of the quinone (Figure 2, Figure S2). This result suggests that the futalosine pathway is functional in *C. trachomatis* and produces MK-7, setting the stage for interrogation of menaquinone biosynthesis as a potential antichlamydial target.

**Figure 2.**
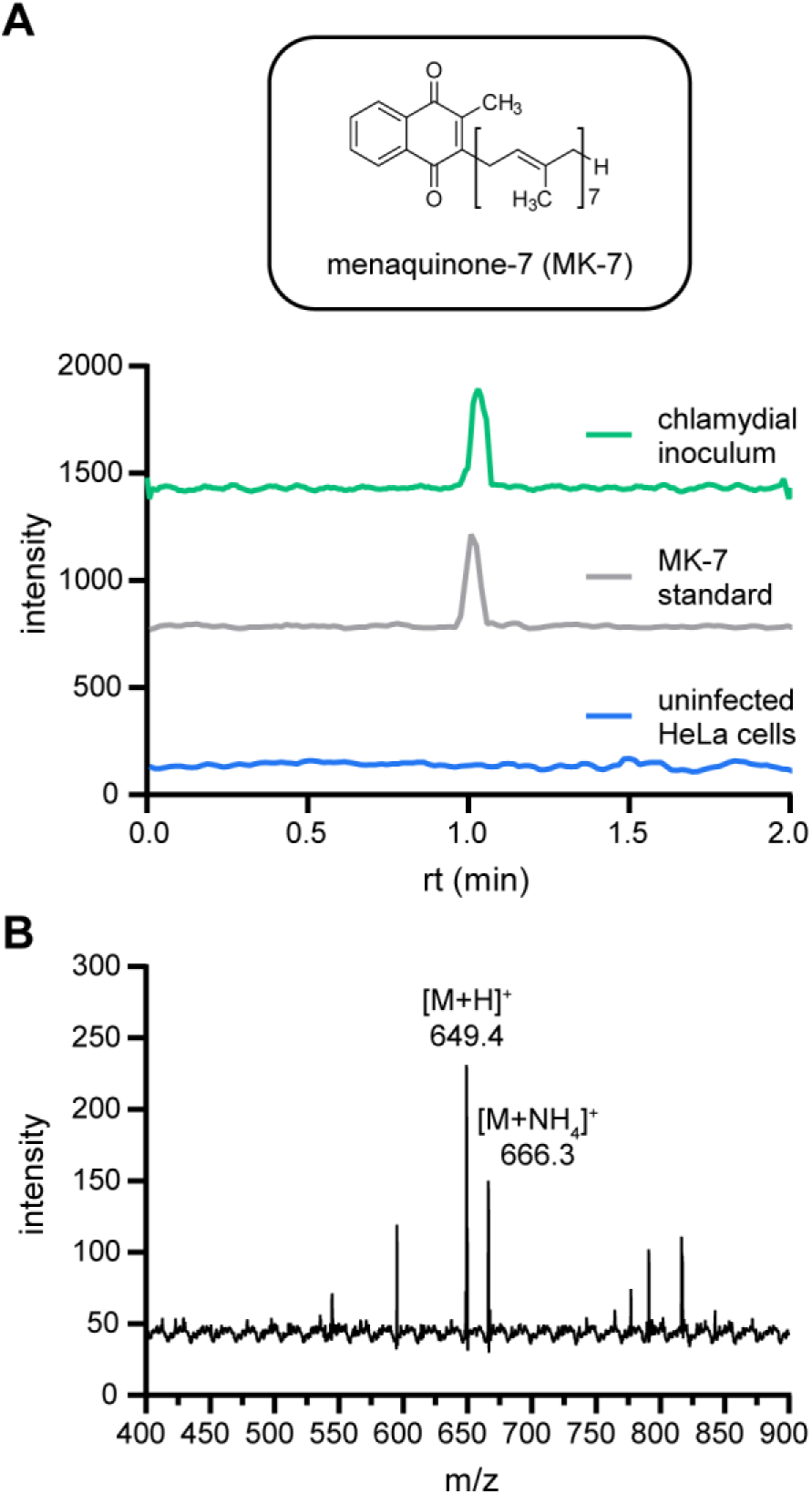
Identification of MK-7 as a chlamydial quinone. **(A)** Extracted ion chromatograms for the m/z 649.4 ([M+H]^+^) to 187.0 fragmentation of MK-7 are shown for chlamydial and HeLa cell extracts as well as an MK-7 standard. **(B)** The mass spectrum for the chlamydial extract contains the [M+H]^+^ and [M+NH4]^+^ ions for MK-7.

### Automated immunofluorescence assay reveals that futalosine pathway inhibitor DHA disrupts inclusion formation

After confirming MK-7 biosynthesis in *C. trachomatis*, we sought to determine the impact of its inhibition on chlamydial infection. A small group of futalosine pathway inhibitors have been discovered previously, including compounds identified through forward^36–39^ and reverse^40^,^41^ chemical genetic approaches. Significantly, several have been found to possess inhibitory activity against the futalosine pathway-containing pathogen *Helicobacter pylori* in mouse models of infection, revealing the translational potential of targeting this pathway.^37^ However, no futalosine pathway inhibitors have been tested for antichlamydial activity to date.

A structurally diverse subset of futalosine pathway inhibitors was chosen to evaluate against *C. trachomatis*: immucillin analogue BuT-DADMe-ImmA,^42^ lasso peptide siamycin I,^37^ and fatty acids 12-methyltetradecanoic acid^36^ and DHA (Figure 3A).^37^ BuT-DADMe-ImmA, the only inhibitor of the group with a characterized target within the futalosine pathway, is a 5’-methylthioadenosine nucleosidase (MTAN) transition state analogue and an inhibitor of MqnB.^42^ The remaining three inhibitors were identified in disk diffusion screens, where each compound inhibited the growth of a futalosine pathway-containing bacterium (*Bacillus halodurans* C-125 or *Kitasatospora setae* KM-6054), but not a classical menaquinone biosynthesis pathwaycontaining strain *(Bacillus subtilis* H17).^37^ While their specific molecular targets within the futalosine pathway have yet to be identified, several mechanistic hypotheses have been formulated. Both 12-methyltetradecanoic acid and DHA are hypothesized to inhibit the membrane-bound prenyltranferase MqnP based on 1) structural similarity to its polyisoprene pyrophosphate substrate and 2) a previous study which suggested that free fatty acids inhibit a step after the formation of futalosine.^36^ Siamycin I has been recently characterized as a binder of the peptidoglycan biosynthesis intermediate lipid II, which contains an undecaprenyl pyrophosphoryl motif;^43^ therefore, it is likely that siamycin I interacts with the MqnP substrate heptaprenyl pyrophosphate, given its structural similarity to lipid II.

**Figure 3.**
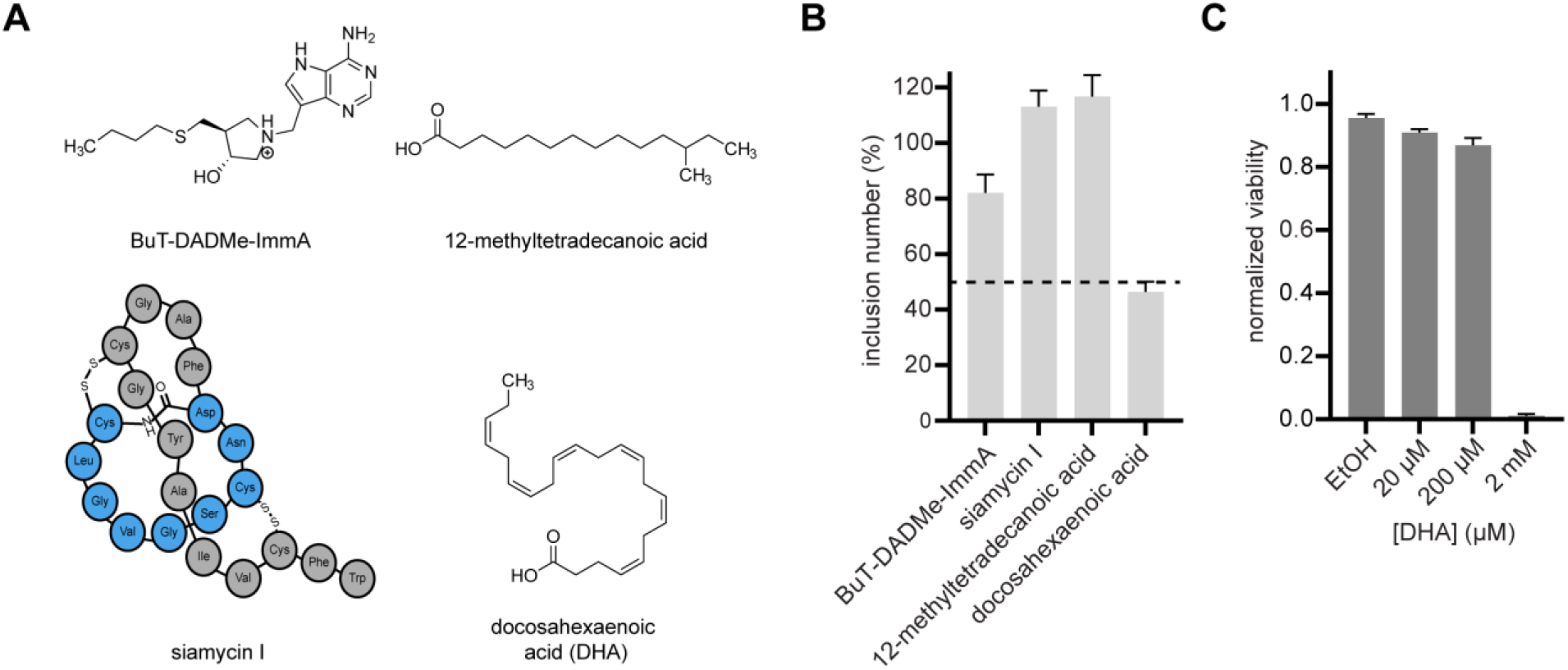
An automated immunofluorescence assay monitoring inclusion formation in the presence of futalosine pathway inhibitors reveals DHA as an antichlamydial agent. **(A)** Structures of the previously identified futalosine pathway inhibitors tested against *C. trachomatis* in this study. **(B)** *C. trachomatis-infected* HeLa cells were treated with inhibitors to characterize their effects on inclusion formation. Inclusion number was quantified using the CellInsight CX5 High-Content Screening Platform. Data are visualized as the inclusion number from each inhibitor treatment relative to its respective vehicle control (DMSO for BuT-DADMe-ImmA and siamycin I; ethanol for 12-methyltetradecanoic acid and DHA) and given as the mean of triplicate samples ± s.e.m. As the only compound that reduced inclusion number by >50%, DHA was selected as a hit. **(C)** MTT cell viability assay confirms that DHA treatment results in >85% HeLa cell viability up to 200 μM. Data are visualized as viability normalized to an untreated control (mean of triplicate samples ± s.e.m).

To test the antichlamydial effects of these inhibitors, we developed an automated immunofluorescence-based assay of inclusion formation in a HeLa cell model of chlamydial infection. HeLa cells were infected with *C. trachomatis* at a multiplicity of infection (MOI) of 0.5, then treated with each inhibitor (125 μM) or vehicle control for 40 hpi. Immunofluorescence staining for the chlamydial inclusion membrane was performed, and a high-content imaging platform was implemented to monitor inclusion formation, with the goal of identifying a compound that inhibited inclusion formation by >50% relative to its respective vehicle control. Using this assay, one futalosine pathway inhibitor, DHA, was discovered to meet this threshold (Figure 3B). The remaining three inhibitors caused little to no decrease in inclusion number, which we hypothesize may be attributed to a lack of permeability. Because of the intracellular nature of *C. trachomatis*, cell entry is a well-established barrier to the efficacy of antichlamydial therapeutics.^44^ Furthermore, difficulties with cellular uptake have been previously demonstrated for siamycin I^43^ and immucillin-type transition state analogues such as BuT-DADMe-ImmA.^45,46^

Despite these permeability challenges, the decrease in inclusion number caused by treatment of *C. trachomatis*-infected HeLa cells with 125 μM DHA is, to our knowledge, the first time a futalosine pathway inhibitor has been found to disrupt chlamydial infection. Notably, the viability of HeLa cells was not compromised at the tested DHA concentrations as measured with a 3-(4,5-dimethylthiazol-2-yl)-2,5-diphenyl tetrazolium bromide (MTT) assay (>85% viability at 200 μM DHA), indicating that the observed activity of DHA is not attributed to host cell toxicity (Figure 3C).

### DHA inhibits multiple aspects of chlamydial infection

To expand our analysis of the antichlamydial effects of DHA to several characteristics of chlamydial infection, *C. trachomatis*-infected HeLa cells were challenged with a concentration gradient of DHA, and the effects of treatment on inclusion number, inclusion size, and infectious progeny were determined. These three parameters were selected to shed light on different elements of the life cycle, as inclusion number, inclusion size, and infectious progeny are indicators of inclusion formation, chlamydial growth, and progression through the life cycle, respectively.

First, inclusion number and size were quantified using the high-content imaging platform previously described after DHA treatment for 40 hpi. A concentrationdependent reduction in inclusion formation was observed, including 83.0% inhibition at 125 μM DHA (Figure 4A, 4B). In contrast, although DHA treatment caused a 28.0% reduction in inclusion number at the lowest tested concentration (31.3 μM), inclusion size was exclusively affected at 125 μM, with a 40.6% decrease relative to vehicle control observed (Figure 4A, 4C). These results reveal that DHA begins to disrupt inclusion formation at lower concentrations but requires a higher concentration to inhibit chlamydial growth. Similar differential effects on inclusion number and size have been previously demonstrated for other antichlamydials such as pyocyanin.^47^

**Figure 4.**
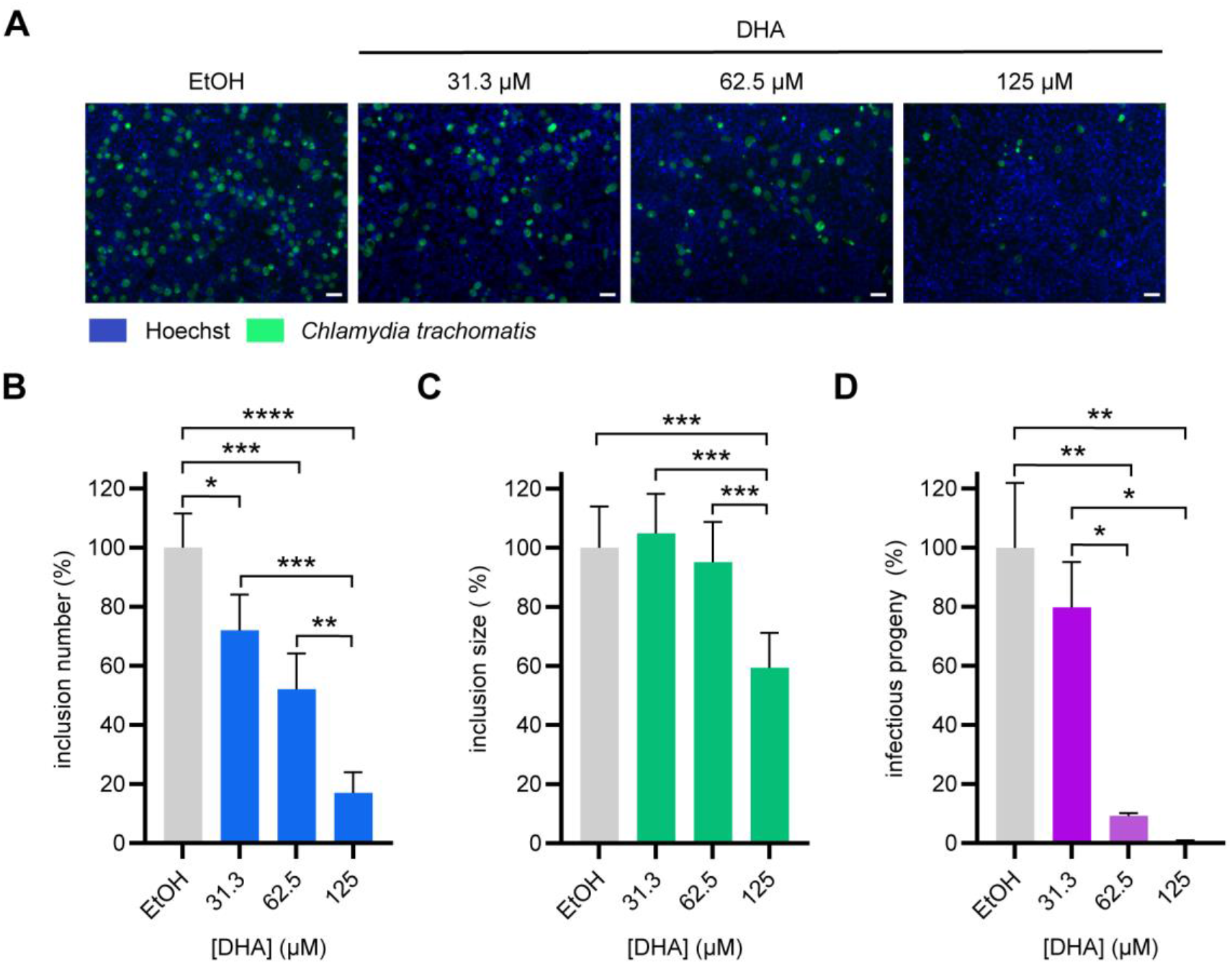
DHA treatment induces several antichlamydial effects. **(A)** Representative immunofluorescence microscopy images for each DHA concentration and vehicle control from the inclusion number experiment are shown. Scale bars represent 50 μm. **(B-D)** [DHA]-dependent effects on inclusion number **(B)**, inclusion size **(C)**, and infectious progeny **(D)** were quantified using the CellInsight CX5 High-Content Screening Platform. Data are visualized as the inclusion number, inclusion size, or infectious progeny from each condition relative to the ethanol vehicle control (mean of 3 biological replicates ± s.e.m.). Significance was determined by RM-ANOVA with Tukey’s post hoc test for pairwise comparisons using JMP,^33^ where **** = p < 0.0001, *** = p < 0.001, ** = p < 0.01, and * = p < 0.05.

To measure the generation of infectious progeny, *C. trachomatis*-infected HeLa cells were harvested after treatment with DHA for 40 hpi, and a titer assay was performed to count recoverable inclusion-forming units (IFUs) from each isolated sample. The effect of DHA on infectious progeny was more consistent with its effect on inclusion number than inclusion size, with a concentrationdependent reduction in progeny observed (Figure 4D). Notably, progeny IFUs were drastically reduced to 0.579% of control IFUs upon treatment with 125 μM DHA. This suggests that DHA impedes the progression of the chlamydial developmental cycle. Collectively, these results establish DHA as a previously unidentified antichlamydial compound that disrupts multiple characteristics of infection.

### Menaquinone-7 rescues the effect of DHA on inclusion formation

Finally, we sought to confirm that the antichlamydial activity of DHA is mediated through inhibition of the futalosine pathway. As part of the initial identification of DHA as a futalosine pathway inhibitor by Yamamato and co-workers, broth cultures of *H. pylori* were treated with DHA alone or in combination with menaquinone-4 (MK-4).^37^ While DHA alone disrupted the growth of *H. pylori*, the inhibition was rescued by MK-4 treatment, validating that DHA targets the futalosine pathway. Inspired by this finding, we hypothesized that defects in *C. trachomatis* infection caused by DHA treatment could be rescued by supplementation with MK-7.

To interrogate this hypothesis, *C. trachomatis-infected* HeLa cells were treated with 125 μM DHA in the presence and absence of 10 μM MK-7 nanoparticles and evaluated for effects on inclusion number, inclusion size, and progeny. MK-7 nanoparticles were developed (Figure S3A, S3B) and employed in these experiments to solubilize MK-7 in cell-tolerable solvents and promote intracellular delivery.^48^ Excitingly, inclusion formation was rescued by cotreatment with MK-7 nanoparticles (Figure 5A, 5B), as the number of inclusions after combination treatment was significantly increased relative to treatment with DHA alone, but not significantly different than the vehicle control. Further, inclusion number was unchanged when infections were treated with MK-7 nanoparticles alone (Figure S3C). These results convey that DHA hinders inclusion formation through inhibition of the futalosine pathway.

**Figure 5.**
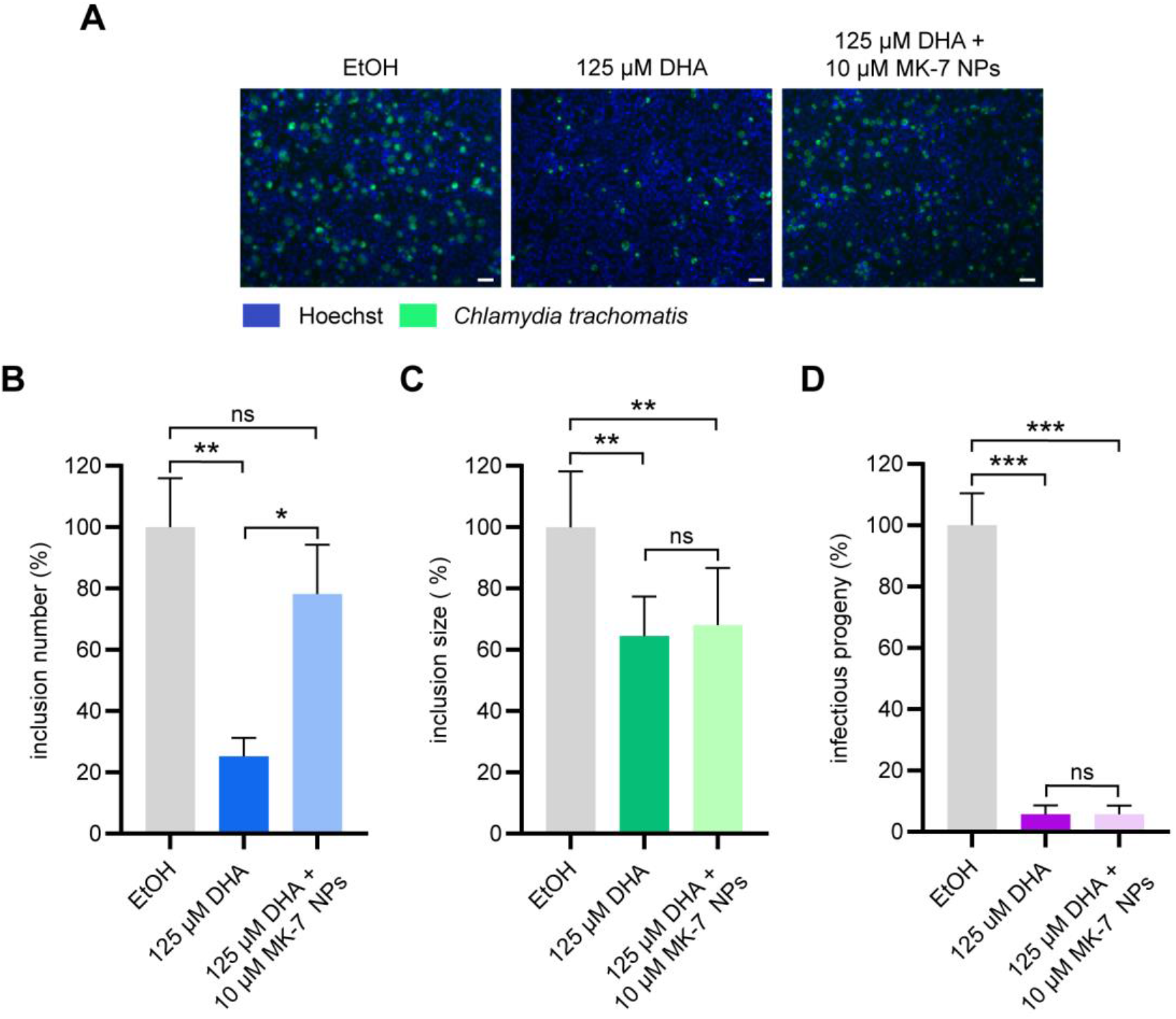
MK-7 nanoparticles rescue DHA-mediated inhibition of inclusion number. **(A)** Representative immunofluorescence microscopy images for each treatment from the inclusion number experiment are shown. Scale bars represent 50 μm. **(B-D)** Inclusion number **(B)**, inclusion size **(C)**, and infectious progeny **(D)** were quantified using the CellInsight CX5 High-Content Screening Platform. Data are visualized as the inclusion number or inclusion size from each condition relative to the ethanol vehicle control (mean of 3 biological replicates ± s.e.m.). Significance was determined by RM-ANOVA with Tukey’s post hoc test for pairwise comparisons using JMP,^33^ where **** = p < 0.0001, *** = p < 0.001, ** = p < 0.01, and * = p < 0.05.

In contrast, inclusion size and infectious progeny were not significantly changed by co-treatment with MK-7 relative to treatment with DHA alone (Figure 5C, 5D). Because polyunsaturated fatty acids have been characterized for their ability to modulate membrane integrity^49^ and inhibit fatty acid synthesis,^50^ we anticipate that DHA may operate through additional mechanisms to affect the various stages of chlamydial infection. Identification of the molecular target(s) of DHA in chlamydial infection is the subject of future work and will enhance our understanding of the differential effects of DHA inhibition and MK-7 rescue. Nonetheless, the rescue of inclusion number by co-treatment with MK-7 is consistent with a mechanism of targeting the futalosine pathway, underscoring the potential of inhibiting menaquinone biosynthesis as an antichlamydial approach.

## CONCLUSIONS

Cumulatively, the results presented herein reveal that the futalosine pathway for menaquinone biosynthesis is a previously unrecognized target for antichlamydial development. The futalosine pathway has been previously hypothesized to play an important role in infection due to the discovery of an active respiratory chain in *C. trachomatis*; however, menaquinone production and the effects of its disruption have remained unexplored. In this work, menaquinone-7 was identified in a chlamydial extract by LC-MS/MS, revealing a functional futalosine pathway. Through an automated immunofluorescence assay of futalosine pathway inhibitors, the polyunsaturated fatty acid DHA was found to inhibit inclusion formation in *C. trachomatis*-infected HeLa cells. DHA was subsequently demonstrated to disrupt both the formation and developmental progression of chlamydial infections in a dose-dependent manner, as 125 μM treatment caused reductions in inclusion number and progeny to 17.0% and 0.579%, respectively, of ethanol vehicle control values. A 40.6% reduction in inclusion size was also observed upon treatment with 125 μM DHA. Significantly, the rescue of inclusion number by co-treatment with 10 μM MK-7 nanoparticles confirms that the mechanism of DHA activity includes inhibition of the futalosine pathway. These results have established for the first time that a futalosine pathway inhibitor disrupts multiple elements of chlamydial infection and have laid the groundwork for inhibitors of this pathway to be explored as novel antichlamydial agents, which are desperately needed to combat this public health threat.

## Acknowledgements

The authors gratefully acknowledge Dr. Raphael Valdivia (Duke University) for the gift of the *Chlamydia trachomatis* L2 strain and Dr. Vern Schramm (Albert Einstein College of Medicine) for the gift of the transition state analogue BuT-DADMe-ImmA used in this work. The authors also acknowledge Dr. Michael Therien (Duke University) and Dr. Mark Wiesner (Duke University) for the use of their respective DLS instruments for nanoparticle measurement. The authors thank Dr. Peter Silinski and the Duke Shared Instrument Facility for mass spectrometry assistance, and Dr. So Young Kim of the Duke Functional Genomics Core Facility for cellomics method development. The authors gratefully acknowledge Dr. Ted Slotkin (Duke University) and Maria Toro Moreno (Duke University) for assistance in the development of statistical analyses and data visualization. Lastly, the authors would like to thank the members of the McCafferty laboratory for thoughtful discourse and feedback throughout the course of this work and preparation of the manuscript.

## Funding and additional information

This work was kindly supported by Duke University, National Institutes of Health Predoctoral Training Grant 5T32GM007105-44 in Pharmacological Sciences to B.M.D., and National Institutes of Health Predoctoral Training Grant 1T32GM133352-01A1 to T.C.O.

## Abbreviations

The abbreviations used are:

ATP: adenosine triphosphate;
DHA: docosahexaenoic acid;
DMEM: Dulbecco’s modified Eagle’s medium;
EIC: extracted ion chromatogram;
FBS: fetal bovine serum;
IFU: inclusion-forming unit;
LC-MS/MS: liquid chromatography-tandem mass spectrometry;
MK-4: menaquinone-4;
MK-7: menaquinone-7;
MOI: multiplicity of infection;
MS: mass spectrometry;
MTAN: 5’-methylthioadenosine nucleosidase;
MTT: 3-(4,5-dimethylthiazol-2-yl)-2,5-diphenyl tetrazolium bromide;
PBS: phosphate-buffered saline;
s.e.m.: standard error of the mean;
SPG: sucrose-phosphate-glutamate

