## Supplemental Information for "Investigation of the futalosine pathway for menaquinone biosynthesis as a novel target in the inhibition of *Chlamydia trachomatis* infection"

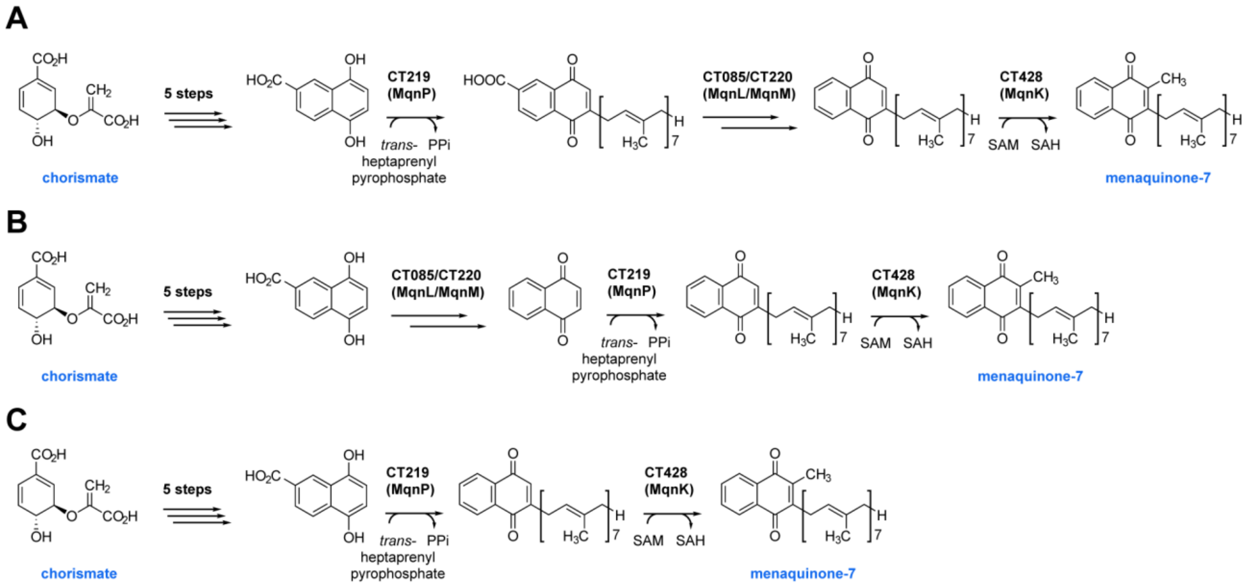


**Figure S1. Potential enzymatic routes to menaquinone-7 in *C. trachomatis*.** As only the first five steps of the futalosine pathway have been characterized to date,[1] the final steps from 1,4-dihydroxy-6-naphthoic acid to menaquinone-7 could occur through several possible orders: **(A)** condensation with heptaprenyl pyrophosphate by the prenyltransferase MqnP, followed by decarboxylation by the UbiX/UbiD-like decarboxylase/flavin prenyltransferase pair MqnL/MqnM, and finally methylation by the SAM-dependent methyltransferase MqnK; **(B)** decarboxylation by MqnL/MqnM, then prenylation by MqnP, and lastly methylation by MqnK; **(C)** prenylation and decarboxylation could occur in a single step catalyzed by MqnP (as occurs in the traditional menaquinone biosynthesis pathway),[2] followed by methylation by MqnK.

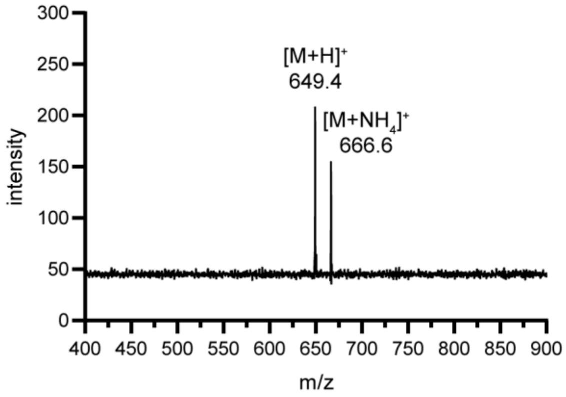


**Figure S2. Mass spectra of commercially acquired menaquinone-7 standard, containing the $[M+H]^+$ and $[M+NH_4]^+$ ions.**

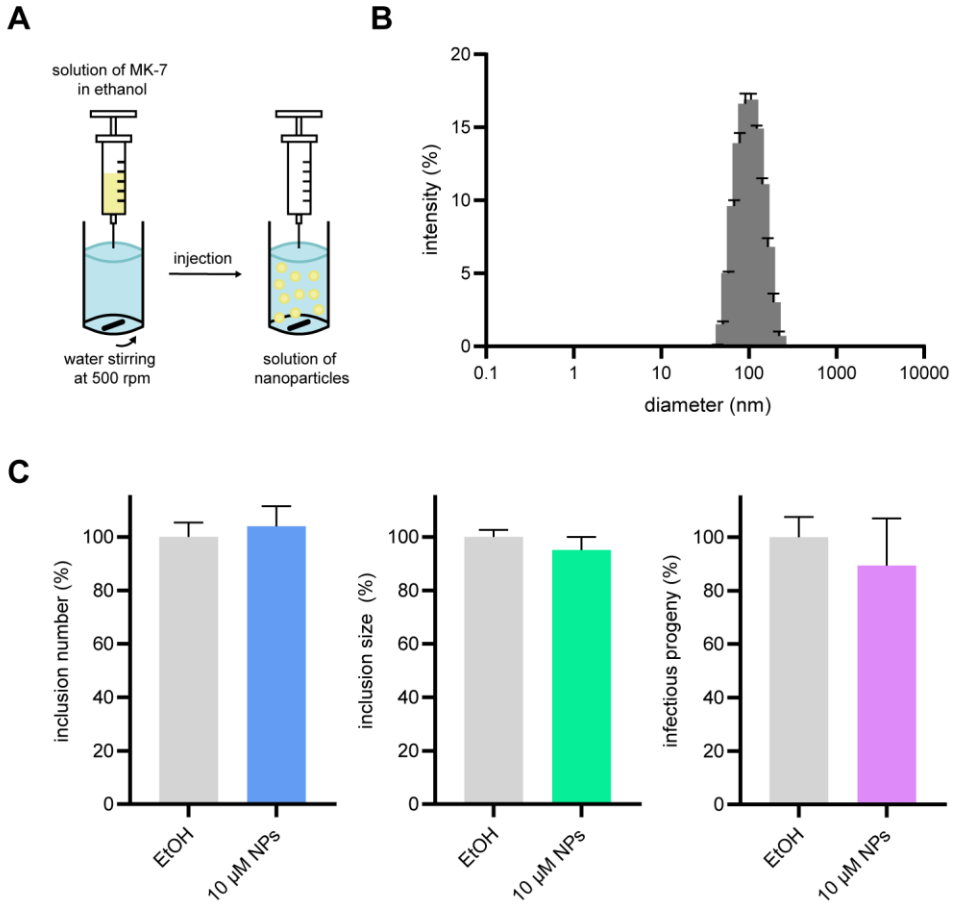


**Figure S3. Production, measurement, and analysis of menaquinone-7 nanoparticles. (A)** Menaquinone-7 nanoparticles were produced by rapid solvent shifting.[3] In this method, a solution of menaquinone-7 in ethanol was rapidly injected into water stirring at 500 rpm, resulting in the spontaneous generation of nanoparticles. **(B)** Diameters of menaquinone-7 nanoparticles were measured by dynamic light scattering (DLS). A representative DLS intensity distribution is shown for a 200 µM menaquinone-7 preparation (average diameter = 106.65 nm). Data are visualized as the mean of four repeat measurements ± standard deviation. **(C)** Menaquinone-7 nanoparticles do not significantly affect chlamydial inclusion number, size, or infectious progeny in a HeLa cell model of infection. Data are visualized as the inclusion number, size, or progeny relative to the ethanol vehicle control and given as the mean of 3 biological replicates ± s.e.m.
